# Automated imaging of duckweed growth and development

**DOI:** 10.1101/2021.07.21.453240

**Authors:** Kevin L. Cox, Jordan Manchego, Blake C. Meyers, Kirk J. Czymmek, Alex Harkess

## Abstract

Duckweeds are some of the smallest angiosperms, possessing a simple body architecture and high rates of biomass accumulation. They can grow near-exponentially via clonal propagation. Understanding their reproductive biology, growth, and development is essential to unlock their potential for phytoremediation, carbon capture, and nutrition. However, there is a lack of non-laborious and convenient methods for spatially and temporally imaging an array of duckweed plants and growth conditions in the same experiment. We developed an automated microscopy approach to record time-lapse images of duckweed plants growing in 12-well cell culture plates. As a proof-of-concept experiment, we grew duckweed on semi-solid media with and without sucrose and monitored its effect on their growth over 3 days. Using the PlantCV toolkit, we quantified the thallus area of individual plantlets over time, and showed that *L. minor* grown on sucrose had an average growth rate four times higher than without sucrose. This method will serve as a blueprint to perform automated high-throughput growth assays for studying the development patterns of duckweeds from different species, genotypes, and conditions.

## Introduction

Duckweeds are aquatic monocots in the Lemnaceae family that comprise the fastest-reproducing land plants (Sree et al., 2015c; Bog et al., 2019). Duckweeds are capable of hyperaccumulating heavy metals and serving as substantial sinks of carbon, nitrogen, and phosphorus (Wang et al., 2016; Ziegler et al., 2017). Furthermore, duckweeds display immense resilience across their global growth range, allowing them to survive in variable temperatures and growth conditions (O’Brien et al., 2020). Duckweeds also serve as a potential source of biofuels and nutritional feed due to their high starch and protein content (Appenroth et al., 2018). Some species are regularly consumed by humans in parts of Southeast Asia (Bhanthumnavin and McGarry, 1971), given its excellent amino acid profile and protein quantity (Appenroth et al., 2018, 2017). Given these unique genetic, growth, and physiological characteristics, in addition to their cosmopolitan global distribution, duckweeds have been proposed as model systems for plant biology (Hillman and Culley, 1978; Cao et al., 2018).

The duckweed family is composed of five genera; *Spirodela*, *Landoltia*, *Lemna, Wolffiella*, and *Wolffia* (An et al., 2018; Bog et al., 2019; Tippery and Les, 2020; Les et al., 2002). Duckweeds lack obvious stems and leaves, instead existing as a leaf-like thallus, and the relative lack of morphological characteristics between species has complicated systematics within the group (Les et al., 1997). Interestingly, each genus has unique features in their growth characteristics and morphology, ranging from the number of roots to their mechanisms of vegetative propagation (Landolt and Kandeler, 1987). Previous microscopy studies have revealed distinct characteristics in duckweed species, such as the high number of stomata, presence/absence of the “pseudoroot”, organization and distribution of chloroplasts, and the location from which new daughter fronds initiate (Sree et al., 2015b; Landolt and Kandeler, 1987; Hoang et al., 2019; Harkess et al., 2021). Notably, since the diameter of these plants ranges from roughly 1-15 mm, it is feasible to image multiple duckweeds simultaneously. For a three-dimensional (3D) perspective, X-ray Computer Tomography (CT) has also been applied to duckweeds, enabling non-destructive imaging of entire plantlets into 3D volumes (Jones et al., 2020). As such, microscopy can play an important role in future developmental research that involves either studies of cell development genes or phenotyping of duckweed species with different genetic backgrounds. However, there is a lack of automated approaches for efficient time-lapse imaging and analysis of duckweed plants under multiple, simultaneous experimental conditions.

Duckweeds are ideal plants to phenotype using automated time-course imaging for several reasons. First, duckweeds are thin plants that can be grown either floating on liquid or a stabilized surface using solid media. Solid media is particularly useful since the plants can be relatively immobilized to remain in a constant focal plane and position, as opposed to the tendency to float out-of-frame in aqueous media. Second, duckweeds grow and reproduce dominantly in one plane of movement, spreading out onto the top of whatever surface they grow on. Thus, imaging plants via a top-positioned camera captures almost all of the available biomass needed to estimate plant size. Third, they can be indefinitely propagated as long as they are kept alive, which is straightforward for most species. Duckweeds already grow in highly adverse conditions around the world (e.g. heat stress, salt stress, UV stress, cold stress, wastewater) (Sree et al., 2015a; O’Brien et al., 2020; Liu et al., 2017; Marín et al., 2007; Uysal and Taner, 2010; Jansen et al., 2001; Fourounjian et al., 2019)), and can be easily stored long-term in dark conditions to induce a state of slow growth hibernation (Jacobs, 1947).

Here we describe the development of an automated approach for imaging of duckweed plants. We performed growth time-lapse imaging of multiple duckweed plants in a single experiment through the use of a microscope with motorized stage, focus and automated imaging capabilities. Furthermore, the high-quality images collected through microscopy were segmented and used to calculate the thallus surface area accumulation over time. Overall, this will enable high-throughput, 2D, automated imaging and computational modeling on accumulated biomass and accelerate future cell and developmental biology studies on this family of fast-growing plants.

## Materials and Methods

### Plant growth conditions

An accession of *Lemna minor* 9253 was obtained from the Rutgers Duckweed Stock Cooperative (RDSC; http://www.ruduckweed.org/). Plants were cultured in sterile flasks in axenic conditions in Hoagland’s liquid media (1.6 g/L Hoagland’s No. 2 Basal Salt Mixture; Sigma Aldrich), with or without 0.5% sucrose, in a growth chamber with the following conditions: constant 22C with a 12 hour photoperiod, 50% relative humidity. Fresh subcultures were made every three weeks.

### Multi-well plate setup

Cell culture plates (12-well; Corning #351143) containing 2 mL/well of sterile Hoagland’s media with 0.7% agarose supplemented with or without 0.5% sucrose were prepared in a sterile biosafety cabinet. After complete cooling of the media, duckweed plants (two-week old cultures) were aseptically transferred to the solidified media. To prevent heavy condensation from accumulating at the top of the plate during the time-course, the cell culture plate lid was made hydrophilic by treatment of a solution containing 0.05% Triton X-100 (Sigma) in 20% ethanol for 30 seconds before pouring off the excess liquid and air-drying (Brewster, 2003). Treated-lids were placed back onto the cell culture plates without parafilm sealing, and the plates were immediately transferred for imaging. While the short time-course experiments in this manuscript did not require wrapping the sample plate edges with parafilm, it is possible that parafilming may be needed for longer time-lapse series to minimize evaporation.

### Automated time-course imaging

12-well plates with duckweed plantlets were imaged with a PlanNeoFluar Z 1.0X objective lens (zoom 7x) in reflected light mode using a Zeiss Axiocam 512 color CCD camera on a ZEISS Axio Zoom.V16 fluorescence macro microscope (ZEISS, Jena, Germany) with a motorized stage and focus. Images were collected via ZEN Blue 2.6 Professional software (ZEISS, Jena, Germany) in “Time Series” and “Tile” modes with 12 location positions (x,y,z coordinates) imaged every 60 minutes following auto-focus at each location for 67 hours. Each location was exported as a series of TIFF images or AVI files for further evaluation and processing. Other than ambient room light, the sample plate was exposed to light only during auto-focus and acquisition period when the Axio Zoom.V16 microscope was collecting images. Each image was acquired with an ~2-15 ms exposure in color at 4248×2322 pixels with a pixel size of 4.429 um.

### Calculation of thallus area

To calculate the duckweed area over time, Docker v2.3.0.4 and Jupyter Notebooks were used to run PlantCV v3.10.0 and OpenCV v3.4.9. All execution of Computer Vision workflows were performed using the Jupyter Notebooks terminal interface. OpenCV was used to extract individual frames from each video (one with sucrose-supplemented media and one without sucrose-supplemented media), outputting 67 frames per time-lapse series. Bash commands were used to simultaneously create a unique directory for each individual frame while PlantCV analyzed the image and output the data to the new directory. The implementation of PlantCV allowed for mask creation, binary image production and then shape analysis. The frames captured from the time-lapse video of duckweed grown without sucrose-supplemented media were cropped using the PlantCV crop function to eliminate non-plant interference (e.g. well edge, glare) captured in the background of the image sequence. No amount of the duckweed thallus area was lost in this process. The frames captured from the time-lapse series of duckweed grown on sucrose-supplemented media were not cropped at any point in this workflow. Each JSON file output by the PlantCV workflow containing thallus area data in pixels (located in each frame’s output directory) was moved to a new unique directory containing only these JSON files. Using Bash, this directory was sorted by the frame number located at the end of each file name (“frame_1.json”, “frame_2.json” etc). This sorted list was saved as a text file and all JSON files were appended using Python. The three parameters ‘observations,’ ‘area,’ and ‘value’ were specified to extract the number of pixels each duckweed occupied in its image and plotted in R v4.0.4. All code is available on Github at https://github.com/plantbiojordan/biointernship.

## Results

### Development of an automated imaging platform

Our goal was to implement a system for automated imaging and analysis of duckweeds under different growth conditions in an efficient manner. Additionally, this would provide an improved platform to utilize algorithms for calculating plant growth. Therefore, we developed a workflow from sample preparation to quantitative data analysis as illustrated in Figure 1. Wells of a multi-well plate were first filled with semi-solid Hoagland’s media. Semi-solid media was used, instead of liquid media, to reduce the mobility of the duckweeds, allowing the plants to remain in view and with the same orientation during the time-lapse series. After placing one axenic duckweed cluster per well in the 12-well plate, the time-lapse was conducted using a ZEISS Axio Zoom.V16 macro microscope with motorized stage and focus. An advantage of using this microscope was the ability to set up the device to automatically take images with large fields- at specified time points. Another advantage is the motorized stage made it feasible to repeatedly image individual wells of a multi-well plate in an unattended fashion. After completion of the time-lapse, the images were analyzed via PlantCV, an open-source image analysis software package targeted for plant phenotyping (Gehan et al., 2017). Lastly, the data obtained from the computational analyses were plotted on a graph.

**Figure 1.**
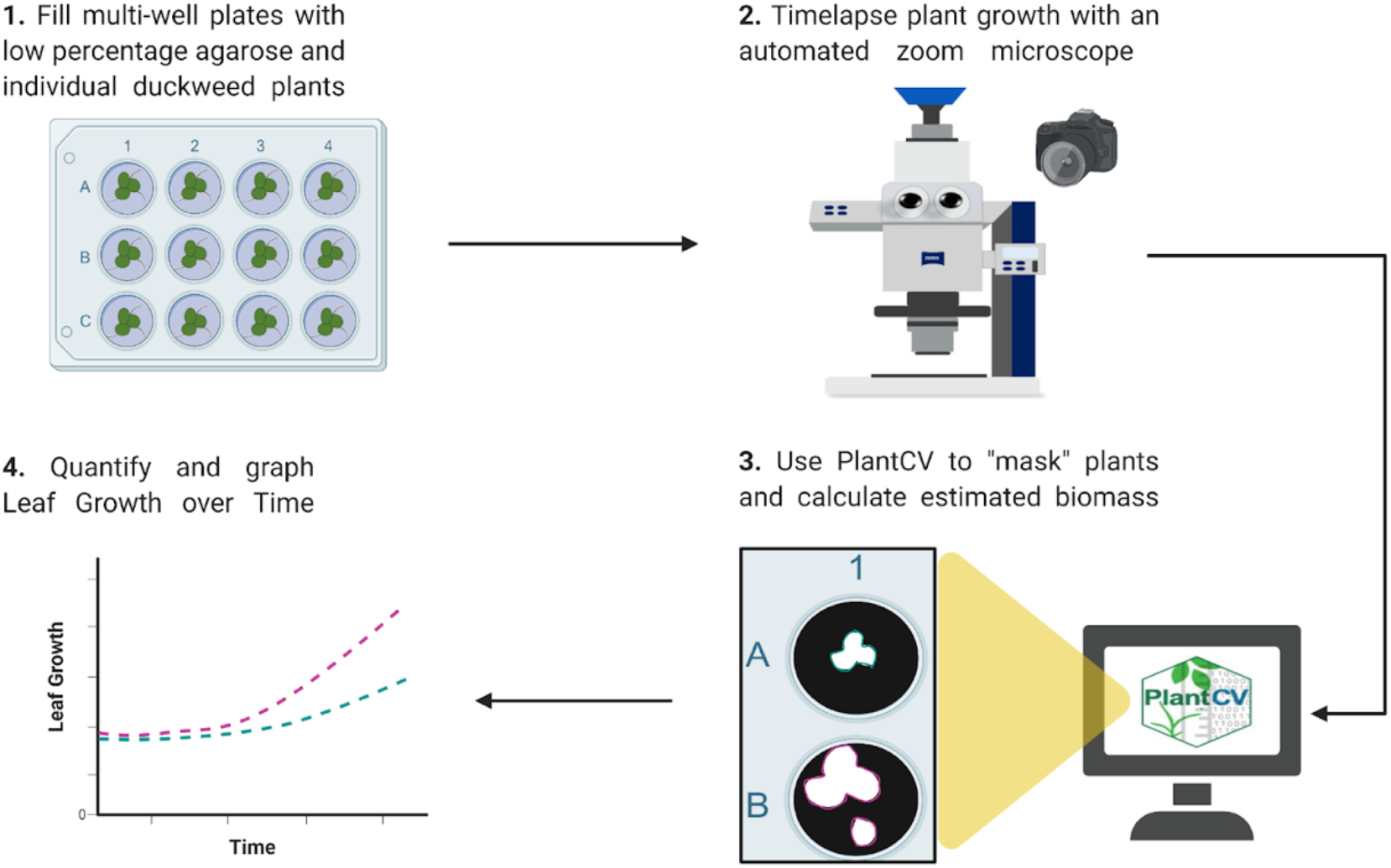
Overview of Automated Imaging and Analysis Platform. Created with BioRender.com.

### Time-lapse imaging of duckweed growth

We conducted a 67 hour growth time-lapse on duckweed accessions *L. minor* 9253. In the 12-well plate, half of the wells were supplemented with 0.5% sucrose. This variable was added to compare the growth rates of the duckweeds growing on sucrose to the ones growing without sucrose. We obtained high quality images of individual duckweeds hourly over the course of 67 hours. As expected, the acquired images revealed that *L. minor* plants growing on media with sucrose grew faster (**Fig. 2, Movie S1**) compared to the plants growing on media without supplemented sucrose (**Fig. 2, Movie S2**).

**Figure 2.**
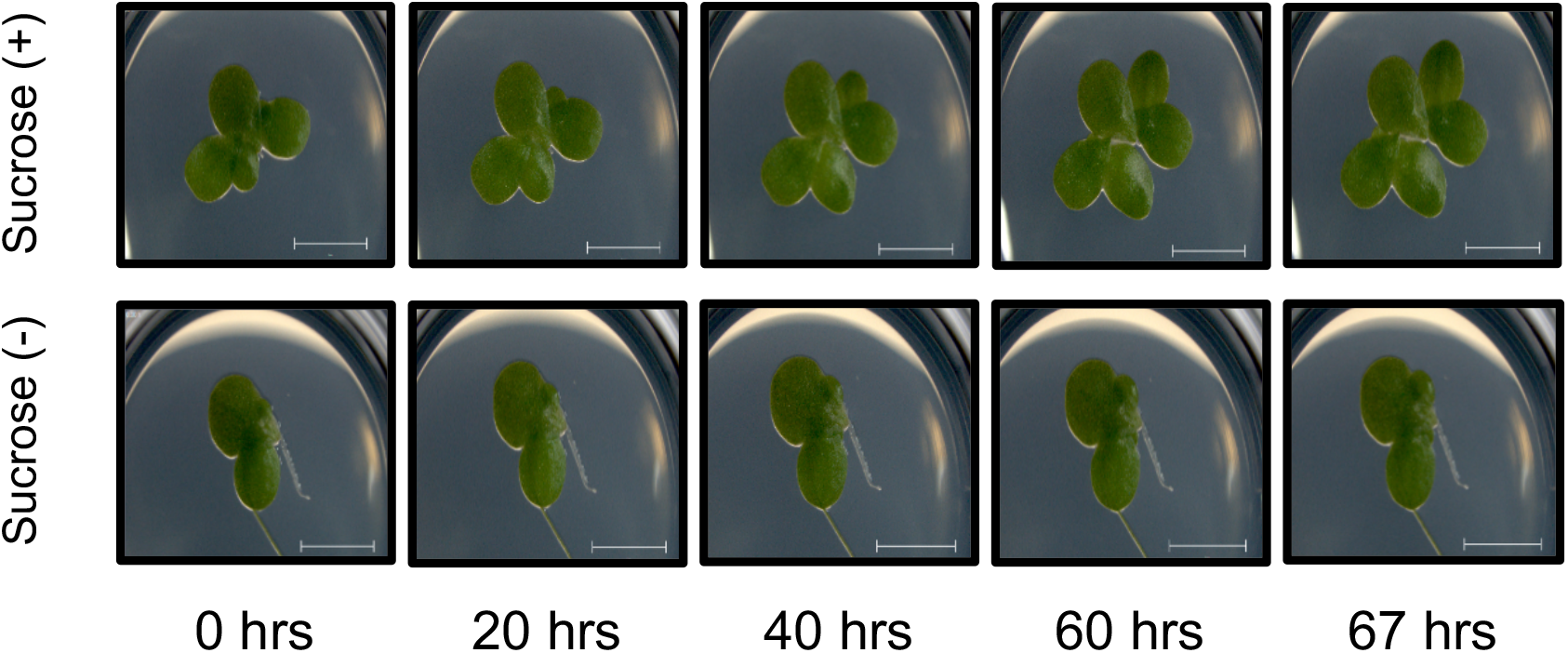
Time-course Imaging of *Lemna minor*. Plants were grown in 12-well plates with solid media with and without sucrose additions. Scale bars: 5 mm.

### Image-based analysis of thallus area

To calculate the thallus area for each of the images acquired by the Axio Zoom.V16 microscope, we developed a computational processing routine using the PlantCV and OpenCV platforms. After extracting the raw individual frames from the time-lapse videos (**Fig. 3A**), the processing routine was designed to identify individual duckweed thalli and measure their areas, lengths, and pixel counts (**Fig. 3**). After completion of the processing routine, the pixel numbers obtained from each of the 67 frames were quantified on a graph to represent the duckweed thallus growth over the time-lapse. The *L. minor* plants growing on media supplemented with sucrose had a nearly 4-times higher average hourly growth rate percent (0.63%) compared to the *L. minor* plants growing on media without sucrose (0.15%) (**Fig. 3D**), supporting our findings in Figure 2.

**Figure 3.**
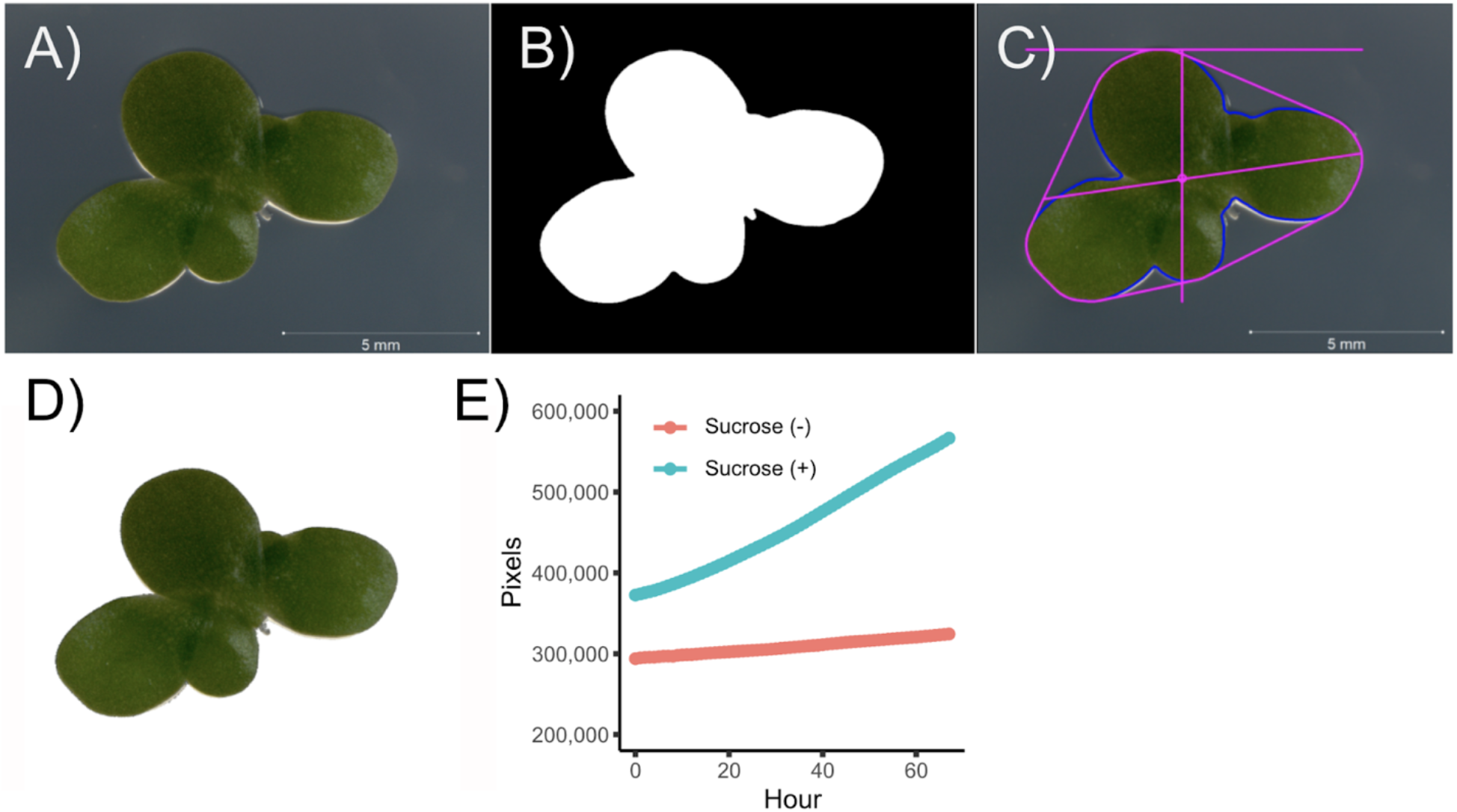
PlantCV phenotyping workflow and thallus area calculation. **(A)** Raw images are acquired from the Axio Zoom.V16. **(B)** Binary image to separate plant from background. **(C)** Object identification, outlining, and shape identification. **(D)** Masked image with background removed. **(E)** Estimates of thallus area growth rate based on pixel area are plotted across the duration of the time-lapse. Red color represents *L. minor* grown without sucrose, while turquoise color represents *L. minor* grown with sucrose.

## Discussion

Duckweeds have high bioremediation and bioenergy feedstock potential (Oron, 1994). The development of platforms to phenotype these plants in a non-laborious, semi high-throughput manner is critical for pairing with genome-scale data to understand which gene families and pathways are involved in duckweed growth and development. Here we describe a protocol to obtain high quality images of duckweed and to serve as a proxy for quantifying their accumulated biomass in our time lapse growth data. This involved the combination of a macro microscope with automated image acquisition capabilities and a computational processing routine for semi high-throughput phenotyping analysis. We applied this workflow in a proof-of-concept experiment that recorded a 67 hour growth time-lapse of *L. minor* growing on media with or without supplemented sucrose and measured and compared their accumulated biomass. With this procedure described in this method, other potential experiments that are now feasible include phenotyping multiple genetic backgrounds and performing a variety of growth assays on a single multi-well plate.

While this is a powerful platform for high resolution, automated imaging, one limitation is the accessibility to the required equipment. In this protocol, we used a stereo and zoom microscope with automated imaging. It may be possible to use other microscopes as alternatives, such as stationary Raspberry Pi-based imaging systems that capture all plants at the same time, but this may increase the number of labor intensive steps and decrease image quality. A secondary, but relatively minor, limitation is that our protocol has duckweeds placed in wells containing semi-solid media instead of liquid media, a state of matter that better simulates their natural growing environment. We found that plants placed in liquid media tended to float out of the field-of-view over the course of the time-lapse. However, our testing of the smallest duckweed species showed that *Wolffia microscopica* tended to “tumble” and roll in semi-solid media, as opposed to staying flat, indicating that species-specific considerations and optimizations need to be taken into account during experimental planning. Nevertheless, this platform will potentially be transformative for future duckweed studies. The application of this platform for semi high-throughput phenotyping will accelerate genetic, molecular, and development studies across all duckweed species.

## Acknowledgments

This work was supported by a Howard Hughes Medical Institute Hanna H. Gray Fellowship to KLC, a HudsonAlpha BioTrain summer internship to JM, and an NSF PGRP Post-doctoral Fellowship to AH (IOS-PGRP #1611853). Additional support for the work comes from the Donald Danforth Plant Science Center, including the Advanced Bioimaging Laboratory (RRID:SCR_018951).

## Authors Contributions

KLC, KJC, and AH conceived the experiments; KLC, JM, KJC, and AH conducted the experiments and analyses; KLC, JM, BCM, KJC, and AH wrote and edited the manuscript.

## References

1. An, D., Li, C., Zhou, Y., Wu, Y., and Wang, W. (2018). Genomes and Transcriptomes of Duckweeds. Front Chem 6: 230.

2. Appenroth, K.-J., Sowjanya Sree, K., Bog, M., Ecker, J., Seeliger, C., Böhm, V., Lorkowski, S., Sommer, K., Vetter, W., Tolzin-Banasch, K., and Others (2018). Nutritional Value of the Duckweed Species of the Genus Wolffia (Lemnaceae) as Human Food. Frontiers in Chemistry 6: 483.

3. Appenroth, K.J., Sree, K.S., Böhm, V., Hammann, S., Vetter, W., Leiterer, M., and Jahreis, G. (2017). Nutritional value of duckweeds (Lemnaceae) as human food. Food Chem. 217: 266–273.

4. Bhanthumnavin, K. and McGarry, M.G. (1971). Wolffia arrhiza as a possible source of inexpensive protein. Nature 232: 495.

5. Bog, M., Appenroth, K.-J., and Sree, K.S. (2019). Duckweed (lemnaceae): Its molecular taxonomy. Front. Sustain. Food Syst. 3: 117.

6. Brewster, J.D. (2003). A simple micro-growth assay for enumerating bacteria. J. Microbiol. Methods 53: 77–86.

7. Cao, H.X., Fourounjian, P., and Wang, W. (2018). The Importance and Potential of Duckweeds as a Model and Crop Plant for Biomass-Based Applications and Beyond. In Handbook of Environmental Materials Management, C.M. Hussain, ed (Springer International Publishing: Cham), pp. 1–16.

8. Fourounjian, P., Tang, J., Tanyolac, B., Feng, Y., Gelfand, B., Kakrana, A., Tu, M., Wakim, C., Meyers, B.C., Ma, J., and Messing, J. (2019). Post-transcriptional adaptation of the aquatic plant Spirodela polyrhiza under stress and hormonal stimuli. Plant J. 98: 1120–1133.

9. Gehan, M.A., Fahlgren, N., Abbasi, A., Berry, J.C., Callen, S.T., Chavez, L., Doust, A.N., Feldman, M.J., Gilbert, K.B., Hodge, J.G., and Others (2017). PlantCV v2: Image analysis software for high-throughput plant phenotyping. PeerJ 5: e4088.

10. Harkess, A., McLoughlin, F., Bilkey, N., Elliott, K., Emenecker, R., Mattoon, E., Miller, K., Czymmek, K., Vierstra, R., Meyers, B.C., and Others (2021). An improved Spirodela polyrhiza genome and proteomic analyses reveal a conserved chromosomal structure with high abundances of chloroplastic proteins favoring energy production. J. Exp. Bot. 72: 2491–2500.

11. Hillman, W.S. and Culley, D.D. (1978). The Uses of Duckweed: The rapid growth, nutritional value, and high biomass productivity of these floating plants suggest their use in water treatment, as feed crops, and in energy-efficient farming. Am. Sci. 66: 442–451.

12. Hoang, P.T.N., Schubert, V., Meister, A., Fuchs, J., and Schubert, I. (2019). Variation in genome size, cell and nucleus volume, chromosome number and rDNA loci among duckweeds. Sci. Rep. 9: 3234.

13. Jacobs, D.L. (1947). An Ecological Life-History of Spirodela Polyrhiza (Greater Duckweed) with Emphasis on the Turion Phase. Ecol. Monogr. 17: 437–469.

14. Jansen, M.A., van den Noort, R.E., Tan, M.Y., Prinsen, E., Lagrimini, L.M., and Thorneley, R.N. (2001). Phenol-oxidizing peroxidases contribute to the protection of plants from ultraviolet radiation stress. Plant Physiol. 126: 1012–1023.

15. Jones, D.H., Atkinson, B.S., Ware, A., Sturrock, C.J., Bishopp, A., and Wells, D.M. (2020). Preparation, Scanning and Analysis of Duckweed Using X-Ray Computed Microtomography. Front. Plant Sci. 11: 617830.

16. Landolt, E. and Kandeler, R. (1987). Biosystematic investigations in the family of duckweeds (Lemnaceae), Vol. 4: The family of Lemnaceae - a monographic study, Vol. 2 (phytochemistry, physiology, application, bibliography). Veroeffentlichungen des Geobotanischen Instituts der ETH, Stiftung Ruebel (Switzerland) 95: 638.

17. Les, D.H., Crawford, D.J., Landolt, E., Gabel, J.D., and Kimball, R.T. (2002). Phylogeny and Systematics of Lemnaceae, the Duckweed Family. Systematic Botany 27: 221–240.

18. Les, D.H., Landolt, E., and Crawford, D.J. (1997). Systematics of the Lemnaceae (duckweeds): Inferences from micromolecular and morphological data. Pl. Syst. Evol. 204: 161–177.

19. Liu, C., Dai, Z., and Sun, H. (2017). Potential of duckweed (Lemna minor) for removal of nitrogen and phosphorus from water under salt stress. J. Environ. Manage. 187: 497–503.

20. Marín, C.M.D.-C., Del-Campo Marín, C.M., and Oron, G. (2007). Boron removal by the duckweed Lemna gibba: A potential method for the remediation of boron-polluted waters. Water Research 41: 4579–4584.

21. O’Brien, A.M., Yu, Z.H., Luo, D.-Y., Laurich, J., Passeport, E., and Frederickson, M.E. (2020). Resilience to multiple stressors in an aquatic plant and its microbiome. Am. J. Bot. 107: 273–285.

22. Oron, G. (1994). Duckweed culture for wastewater renovation and biomass production. Agric. Water Manage. 26: 27–40.

23. Sree, K.S., Adelmann, K., Garcia, C., Lam, E., and Appenroth, K.-J. (2015a). Natural variance in salt tolerance and induction of starch accumulation in duckweeds. Planta 241: 1395–1404.

24. Sree, K.S., Maheshwari, S.C., Boka, K., Khurana, J.P., Keresztes, Á., and Appenroth, K.-J. (2015b). The duckweed Wolffia microscopica: A unique aquatic monocot. Flora - Morphology, Distribution, Functional Ecology of Plants 210: 31–39.

25. Sree, K.S., Sudakaran, S., and Appenroth, K.-J. (2015c). How fast can angiosperms grow? Species and clonal diversity of growth rates in the genus Wolffia (Lemnaceae). Acta Physiol. Plant 37: 204.

26. Tippery, N.P. and Les, D.H. (2020). Tiny Plants with Enormous Potential: Phylogeny and Evolution of Duckweeds. In The Duckweed Genomes, X.H. Cao, P. Fourounjian, and W. Wang, eds (Springer International Publishing: Cham), pp. 19–38.

27. Uysal, Y. and Taner, F. (2010). The Evaluation of the Pb(II) Removal Efficiency of Duckweed Lemna Minor (L.) from Aquatic Mediums at Different Conditions. Survival and Sustainability: 1107–1116.

28. Wang, W., Li, R., Zhu, Q., Tang, X., and Zhao, Q. (2016). Transcriptomic and physiological analysis of common duckweed Lemna minor responses to NH4(+) toxicity. BMC Plant Biol. 16: 92.

29. Ziegler, P., Sree, K.S., and Appenroth, K.J. (2017). The uses of duckweed in relation to water remediation. Desalination Water Treat. 63: 327–342.

